# mLDM: a new hierarchical Bayesian statistical model for sparse microbioal association discovery

**DOI:** 10.1101/042630

**Authors:** Yuqing Yang, Ning Chen, Ting Chen

**Affiliations:** Bioinformatics Division and Center for Synthetic & Systems Biology, TNLIST, China; Department of Computer Science and Technology, Tsinghua University, Beijing 100084, China; State Key Lab of Intelligent Technology and Systems, Tsinghua University, Beijing 100084 China; Program in Computational Biology and Bioinfomatics, University of Southern California, CA 90089 USA

**Author notes:** Corresponding authors: Ting Chen and Ning Chen. This work is supported by the National Natural Science Foundation of China (Nos: 61305066, 61561146396, 61322308).

## Abstract

Interpretive analysis of metagenomic data depends on an understanding of the underlying associations among microbes from metagenomic samples. Although several statistical tools have been developed for metagenomic association studies, they suffer from compositional bias or fail to take into account environmental factors that directly affect the composition of a given microbial community. In this paper, we propose <u>m</u>etagenomic <u>L</u>ognormal-<u>D</u>irichlet-<u>M</u>ultinomial (mLDM), a hierarchical Bayesian model with sparsity constraints to bypass compositional bias and discover new associations among microbes and between microbes and environmental factors. The mLDM model can 1) infer both conditionally dependent associations among microbes and direct associations between microbes and environmental factors; 2) consider both compositional bias and variance of metagenomic data; and 3) estimate absolute abundance for microbes. Thus, conditionally dependent association can capture direct relationship underlying microbial pairs and remove the indirect connections induced from other common factors. Empirical studies show the effectiveness of the mLDM model, using both synthetic data and the TARA Oceans eukaryotic data by comparing it with several state-of-the-art methodologies. Finally, mLDM is applied to western English Channel data and finds some interesting associations.

## 1 Introduction

Understanding interactions among microbes and between microbes and their environment is a key research topic in microbial ecology [28]. Most microbes cannot be cultured in laboratories, making it difficult to gain an understanding of their interactions with existing technologies. However, with the advancement of high-throughput sequencing technology, we are able to sequence 16s rRNA genes or whole metagenome of uncultured microbes directly from samples at diverse time or spots and, as a result, obtain microbial abundance information [47] for further exploration. Various microbial datasets from different environments, such as oceans, soils and humans have been published [41, 6, 37] over the last few years. One of the major challenges is to discover associations, usually referred to as positive and negative relationships, among microbes and between microbes and environmental factors. Such associations could help us to unravel real interactions, including, for example, commensalism, parasitism and competition in a community, resulting in a broad understanding of community-wide dynamics.

Associations can be measured by different statistical methods to show reasonable relationships. Existing association studies can be classified into two categories mainly. First, pairwise association calculation, such as Pearson’s correlation coefficient (PCC) and Spearman’s rank correlation coefficient (SCC), computes the correlation between two species. Local similarity association (LSA) also computes pairwise association, but only works with the time-series data[39]. Second, complex association calculation estimates the relationships between one species and the remaining species and/or environmental factors via regression-based methods [17]. Methods of calculating pairwise association are simple, fast and widely adopted [42, 21, 15, 38, 40, 13], but such methods are not suitable for metagenomic datasets for the following two reasons. First, their calculated values may not indicate real associations because of compositional bias which is introduced during the computation of association using computational methods that assume data are unconstrained, while ignoring dependence among the elements of compositional data [3]. More specifically, since the abundance of each microbe in metagenomic samples is usually normalized as the compositional relative abundance by dividing the total read count of a particular sample. Thus, after normalization, the following example shows that x_i_ is not independent from the rest, regardless of their underlying relationships:

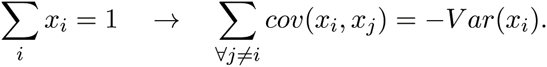

Compositional bias tends to be more severe when some dominant species exist, and is widespread in the marine microbial community [9, 14]. Consequently, for association studies, it is desirable to develop computational methods that bypass compositional bias in order to enable the inference of associations in metagenomic sequencing data. Second, the observed read count of one microbe may deviate from its true abundance based on a given experimental protocol, in which a series of sample preparation, amplification[1], and sequencing steps, leads to large variance of read counts.

Recent advancements have been made in the development of statistical tools to study the associations of data, while taking compositional bias into account. For example, CCREPE [18] estimates the compositionally corrected p-value for every association, which allows the extraction of significant associations via pairwise association calculation. Permutation and bootstrapping are used to generate the null distribution of the association, while considering compositional bias, and the corrected p-value is obtained by the pooled-varianceZ-test. However, the limited number of data samples result in null-distribution and corrected p-values that are unreliable and very sensitive to noises. SparCC [19] infers correlations among microbes by utilizing log-ratio transformation to eliminate the effect of total number of read counts, while imposing sparsity of correlations among microbes.

SPIEC-EASI [29] uses the covariance of the centered log-ratio transformed data to approximate the covariance of log-transformed absolute abundance of microbes and then applies neighborhood selection [33] or standard graphical lasso [20] to obtain the conditionally dependent associations among microbes. However, without considering environmental factors, many associations between and among microbes, as determined by these methods, may not be real. For example, Figure 1 shows that two unrelated microbes (OTU-1 and OTU-2) may appear to be associated just because they both respond to the same environmental perturbation (EF-1). CCLasso [16] is similar to SPIEC-EASI but it estimates the covariance matrix via an alternating direction algorithm instead of the graphical lasso.

**Fig. 1.**
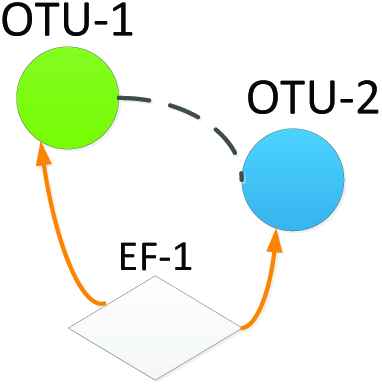
Indirect microbial association. ‘OTU’ and ‘EF’ stand for Operational Taxonomic Units and environmental factor.

Therefore, in this paper, we propose the metagenomic Lognormal-Dirichlet-Multinomial (mLDM) model, a typical hierarchical Bayesian model [2] that learns-complex relationships underlying the data. mLDM could compute conditionally dependent associations among microbes and direct associations between microbes and environmental factors, while takes both compositional bias and variance of metagenomic data into account. In addition, microbial absolute abundance can be estimated, which is useful for further analysis. The effectiveness of mLDM is shown by comparing with the state-of-the-art methods using carefully designed synthetic datasets, and it is further evaluated on TARA Oceans eukaryotic data. Finally, we present the results and findings of mLDM on the TARA Oceans data and western English Channel data.

## 2 Methods

### 2.1 The metagenomic Lognormal-Dirichlet-Multinomial model

Suppose that *N* samples X = 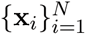. Each x_i_ ϵ ℕ^p^ is a *P*-dimensional vector that contains *P* microbes (or Operational Taxonomic Units (OTUs)), where *x*_*ij*_ represents the sequence/read count of the *j*-th microbes in the i-th sample. Let M = 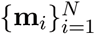 represent the environmental factors, where each m_i_ ϵ ℝ^*Q*^ is a Q-dimensional vector and *m*_*ij*_ represents the value of the *j*-th environmental factor associated with the *i*-th sample.

Figure 2 illustrates the mLDM model for metagenomic sequencing, where x_*i*_ is the read count vector of the *i*-th sample and m_*i*_ records values of the environmental factors corresponding to the *i*-th sample. The latent variable hi is the vector of the relative abundance levels of *P* microbes in the extracted sample, and α_*i*_ represents the absolute abundance levels of the microbes in the original community. We assume that the counts x_*i*_ are proportional to the latent microbial ratios hi which are determined by their absolute abundance α_*i*_. Microbial absolute abundance α_*i*_ can be influenced by two factors: 1) environmental factors m_i_, whose effects on the microbes are denoted by a linear regression model *B*^*T*^ m_*i*_, and 2) the associations among microbes encoded by a latent vector z_i_, which is determined by the matrix *Θ* that records microbial associations and the mean vector *B*_*o*_ that affects the basic absolute abundance of microbes. More specifically, the generative process of the metagenomic Lognormal-Dirichlet-Multinomial hierarchical model is defined as:

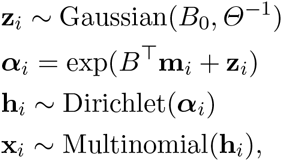

where *B* is a *Q* × *P* parameter matrix, *B*_0_ is the P-dimensional vector, and *Θ* is the inverse covariance matrix (i.e., precision matrix) of a multivariate Gaussian distribution. With this model, our goal is to infer both *B*, the environmental factor-microbe (or EF-OTU) associations, and *Θ*, the microbe-microbe (or OTU-OTU) associations, under some sparsity regularization as will be made clear in next section. We now explain the design of each component in the mLDM model.

**Fig. 2.**
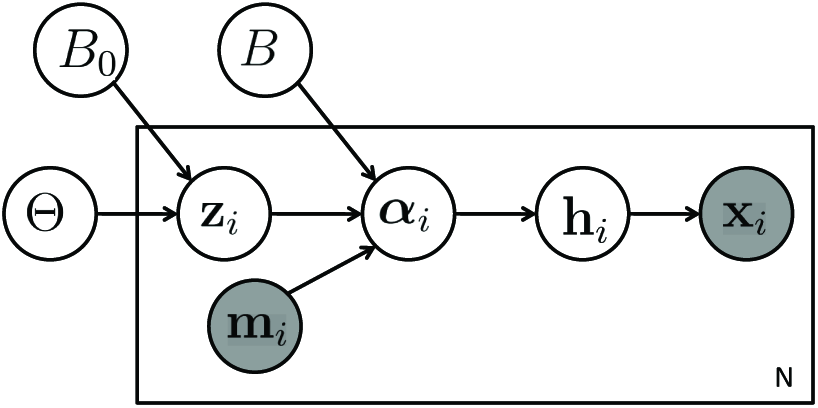
The metagenomic Lognormal-Dirichlet-Multinomial model.

We assume that read count data x_*i*_ follows a multinomial distribution with the microbial ratio parameter h_*i*_:

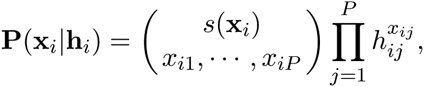

where s(X_*i*_ = 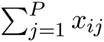 is the total read count of the *i*-th sample. Since the multinomial parameter h_*i*_ is subject to the constraint that 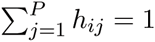, we assume it follows a Dirichlet distribution

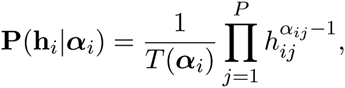

where 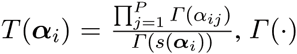 is the Gamma function and 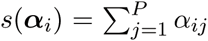. Based on the conjugacy of Dirichlet and multinomial distribution, we can obtain the following Dirichlet-multinomial distribution via integrating h_*ij*_ out

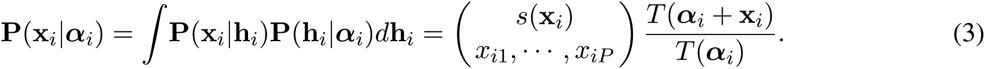

The flexible variance-covariance property of Dirichlet-multinomial distribution is suitable for modeling the sequencing data as mentioned in [44]. A simple explanation follows. We calculate the variance of the real count *x*_*ij*_, **Var***(x_ij_) =* s(X_*i*_). C. *r*_*ij*_. (1 - *r*_*ij*_), and the covariance of two real counts *x*_*ij*_ and *x*_*ik*_, **Cov**(x_*ij*_, *x_ik_)* =-s (X_*i*_). C. *r*_*ij*_. *r*_*ik*_, where C = 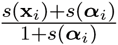 and r_*ij*_ = **α**_*ij*_/s(**α**_*i*_),r_*ik*_ = **α**_*ik*_/s(**α**_*i*_) are true relative abundance levels. We can see that both the variance and covariance of microbial counts are regulated by the sequencing depth s(X_*i*_) and the true relative abundance r_ij_ of the microbes. Moreover, the coefficient between *x*_*ij*_ and *x*_*ik*_ is negative, which models the compositional negative bias.

We further assume that the absolute abundance **α**_*i*_ for all microbes in the i-th sample follows the multivariate lognormal distribution with mean **μ**_*i*_ and covariance *Θ^−1^* which is commonly used to model most microbial abundance except for some occasional species [46, 43, 32]. Microbes survive in a community through conditionally dependent associations. However, at the same time, microbes are also subjected to unpredictable fluctuations impacting their microenvironment. Therefore, we record associations among microbes in the matrix *Θ* and let the mean **μ**_*i*_ vary with the environmental data vector m_*i*_ by a linear regression model. Then the prior distribution is defined as

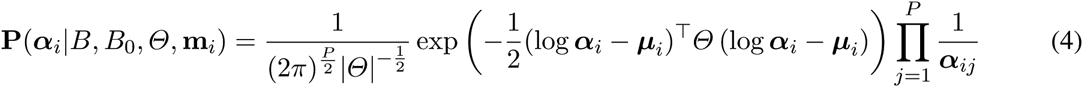

where **μ**_*i*_ = *B*^*T*^ m_*i*_ + *B*_0_. Using the relationship between the lognormal and Gaussian distributions, Eq. (4) is also equivalent to the following form:

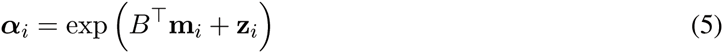

where **Z**_*i*_ ~ *N(B*_0_,Θ^−1^). The formulation in Eq. (5) avoids positivity constraint in the lognormal distribution. This is beneficial for finding the estimates, e.g., by using some unconstrained optimization algorithms, as explained in the next section.

With the above model, we capture both the conditionally dependent associations among microbes and the direct associations between microbes and environmental factors. More specifically, the conditionally dependent associations among microbes are encoded in the precision matrix *Θ.* To visualize the microbial association network, we use an undirected graph denoted as *G*^(1)^ = (*V*^(1)^, *E*^(1)^) employed in the Gaussian Markov random field [35] to represent *Θ*, where V^(1)^ represents the set of nodes denoting *P* microbes and *E*^(1)^ is the set of conditionally dependent associations with each element 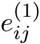 representing the association between the i-th and j-th microbes. If *Θ_ij_ =* 0, then the i-th and the j-th microbes are conditionally independent, and hence, no edge exists between the two microbes in graph *G*^(1)^. The weight of edge 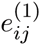, 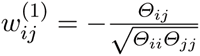, is the strength of the association between the two microbes.

The direct associations between microbes and environmental factors are encoded in weight matrix *B.* The association between the i-th microbe and the j-th environmental factor is *B*_*ji*_, and we can plot them in another bipartite graph *G*^(2)^ = (*V*^(2)^, *E*^(2)^), where the set of nodes *V^(2)^* represents both *P* microbes and *Q* environmental factors, and the edge 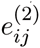 in *E*^(2)^ represents the direct association between the j-th environmental factor and the i-th microbe. The weight of edge 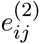 equals 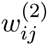 = B_*ji*_.

Overall, our metagenomic association network consists of these two graphs *G*^(1)^ and *G*^(2)^, as illustrated in Figure 3.

**Fig. 3.**
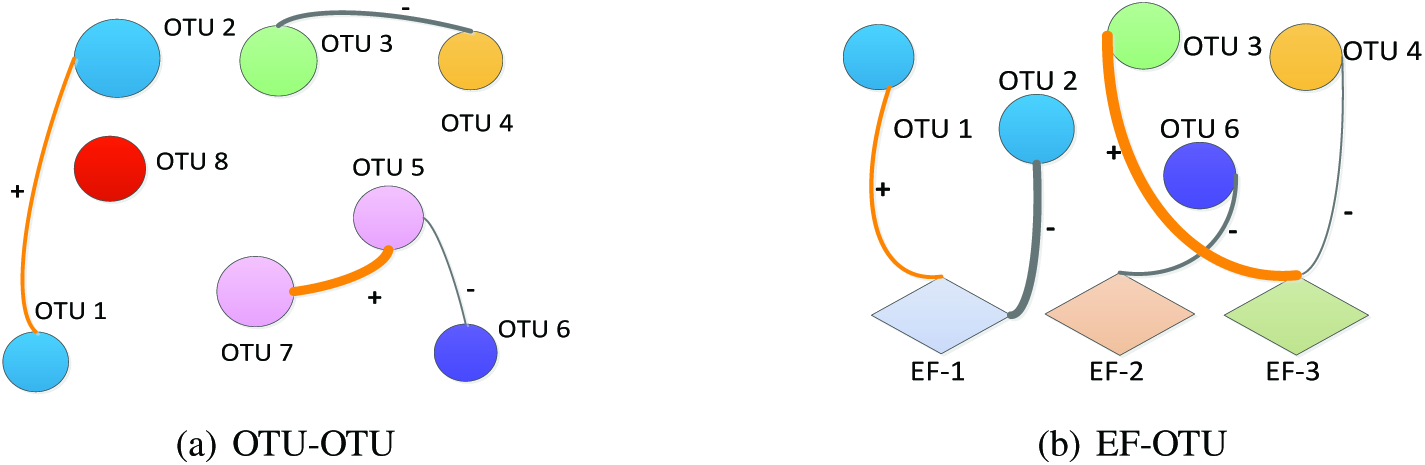
A metagenomic association network is composed of two graphs (a) and (b). ‘+’ and ‘-’ show the positive (orange edge) and negative (grey edge) associations, respectively. (a) is a microbial (OTU-OTU) association graph. (b) is an environmental factor-microbe (EF-OTU) association graph.

### 2.2 Sparse association estimation

We now explain how to estimate the metagenomic association network by using sparsity regularization. Given metagenomic data **X** and environmental factors **M**, the posterior distribution of the latent factors **Z** is

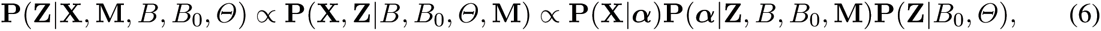

where **P(X**|**α**) can be calculated with Eq. (3), and **P(Z**|B_0_,Θ) *= 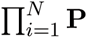* **P**(**Z**_*i*_|B_0_,Θ) with each factor **P**(**Z**_*i*_|B_0_,Θ) being a Gaussian distribution. As a consequence of the deterministic relationship **α**_*i*_ = exp(B^T^m_*i*_ + **Z**_*i*_), it should be noted that the distribution **P**(**α**|**Z**, *B, B*_0_, **M**) is a Dirac delta function. In general, associations among microbes are not expected to be dense and only a few environmental factors will predominate. This motivated us to identify a sparse association network which could be effectively achieved by sparse learning techniques [45]. Also, in practice, the number of samples is usually smaller than the number of microbes, or *N ≪ P.* Therefore, introducing sparsity regularization helps avoid overfitting. Specifically, we estimate the sparse association network by solving the following problem:

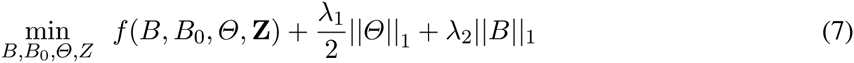

where 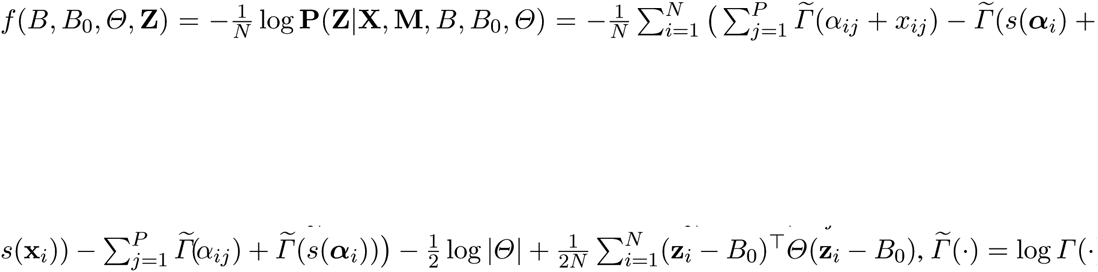 is the log gamma function, and the positive parameters λ_1_ and λ2 are used to control the sparsity of the solution with larger values representing sparser results. Then, the model parameters can be estimated by optimizing the objective function with respect to *Z, B, B*_0_ and Θ alternately.

1. **For Z**, we minimize the objective function in Eq. (7) with respect to Z. Because of independence, we can solve for each **Z**_*i*_ independently by the gradient descent methods. Here, we adopt the limited-memory quasi-Newton (L-BFGS) algorithm [31], which is a quasi-Newton method and converges fast. L-BFGS requires the derivative of *z*_*ij*_, which is computed as follows:

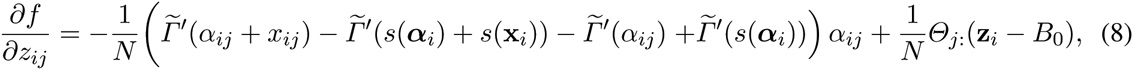

where 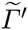*(**α**_ij_)* is the digamma function and *Θ*_j:_ is the jth row of the matrix *Θ*.
2. **For** *B*, we minimize Eq. (7) with respect to *B.* The objective is not differentiable by the existence of the *L1* norm regularizer. Therefore we use the orthant-wise limited-memory quasi-Newton (OWL-QN) algorithm [5], which is based on L-BFGS and can minimize the log likelihood function with *L1* regularization for optimization. The derivative of *B*_*ij*_ is

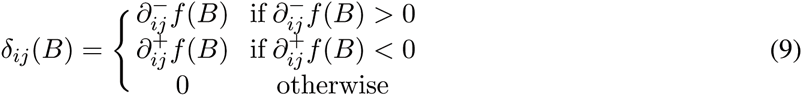

where

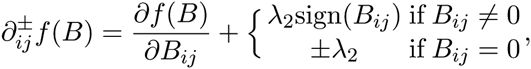

and 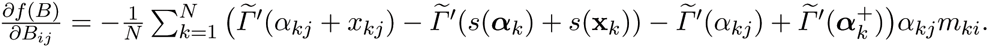.
3. **For** *B*_0_, we have the update rule 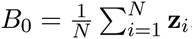, which is the mean of the latent vectors **Z**_*i*_.
4. **For** *Θ*, this step is equal to solving the classical problem of a graphical lasso (glasso):

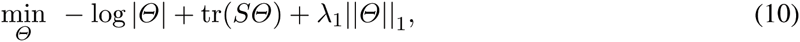

where the empirical covariance *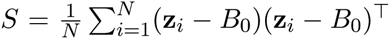*. This problem is also termed as sparse inverse covariance estimation and can be solved with a standard graphical lasso (glasso) algorithm by [20]. However, different from the fully observed glasso, where the empirical covariance is computed once, we should note that our *S* depends on the inferred latent vectors z and needs to update at each iteration. Since z and m mutually influence each other in explaining the observed data x (see the generative process), the learned sparse graph (i.e.,Θ) is affected by environmental factors, matching our intuition in Figure 1.

For model selection, we choose the best parameters for λ1 and λ2 via extended Bayesian information criteria (EBIC) [11]. EBIC improves the original BIC by assigning larger prior to lower dimension models, a strategy more suitable for model selection in large model spaces.

## 3 Results

### 3.1 Synthetic Experiment

To show the effectiveness of the proposed mLDM model, we conducted several experiments and compared mLDM with several state-of-the-art models, including eight programs: PCC, SCC, CCREPE, SparCC, C-CLasso, glasso (graphical lasso), SPIEC (ml) and SPIEC (gl). SPIEC (ml) and SPIEC (gl) are two different modules within SPIEC-EASI, which estimate associations via neighborhood and covariance selection respectively. The first five methods estimate associations via the calculation of correlations with PCC as the baseline, and the last three compute the conditional independence with glasso as the baseline. It should be noted that the Poisson-multivariate normal hierarchical model [7] is not included as a result of its instability when processing high dimensional data. The LSA is also excluded because it requires time series information, which our synthetic dataset doesn’t provide. In the next experiment, we will estimate the following: 1) OTU-OTU associations among all microbes (or OTUs) and 2) EF-OTU associations between environmental factors and microbes.

#### Data Generation Process

The synthetic data can be naturally produced via our generative process. First, the environmental factor matrix M is sampled from the multivariate normal distribution *N*(**0, I**) and then normalized with 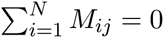 and 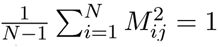. The element *B*_ij_ of matrix *B* is sampled from the uniform distribution of [-0.5, 0.5] and set to 0 with probability of 0.85. Since dominant microbes are found in some microbial communities, we produce vector *B*_0_ by uniformly sampling from [6, 8] with probability of 0.2 and [2,4.5] with probability of 0.8 to affect the distribution of absolute abundance of microbes. To evaluate the ability of mLDM to recover network structures, we follow [29] and use five different precision matrices *Θ* whose adjacency matrices are as follows:

- **Random Graph**: Edge 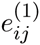 in *E*^(1)^ is set to nonzero with probability 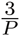 and about 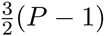 edges are produced.
- **Cluster Graph**: Nodes *V*^(1)^ are randomly split into ⌊P/20⌋ groups and within the same group the nodes *i* and *j* are connected with probability of 0.3.
- **Scale-free Graph**: The B-A algorithm [4] is used to produce a graph in which a) initially two nodes in *G*^(1)^ are connected and b) every new node is added in by linking to a node in the current graph with probability proportional to the degree of the node.
- **Hub Graph**: Nodes *V*^(1)^ are randomly split into ⌊*P*/20⌋ groups, and within the same group, every node is connected with a center node with probability of 1. Finally, random *P* - ⌊*P*/20⌋ edges are included in the *E*^(1)^.
- **Band Graph**: Each adjacent node pair *i* and *j* in *V*^(1)^ is connected if |*i - j* | = 1 and *P* - 1 edges are generated in *E*^(1)^.

We use the huge package [48] to generate *Θ* and obtain the positive definite covariance matrix *Σ = Θ^−1^*. In order to make the covariance matrix *Σ* sparse, and thus beneficial to methods estimating the correlations, we set *Σ*_*ij*_ = 0 if |*Σ_ij_|<* 0.1. Then, **Z**_*i*_ is sampled from the normal distribution *N(0, Σ)*, and **α**_*i*_ is calculated via Equation (5). Next, we generate the Dirichlet-multinomial samples X_*i*_ from Eq. (3). This process relies on the R package “HMP”, which includes the generation of Dirichlet-multinomial samplers. For *B, B*_0_ and *Θ* with five structures, all methods are compared with the following four experimental settings: *P =* 50, *Q =* 5 and *N =* 25, 50,200, and 500. We use public codes glasso, CCREPE, SPIEC-EASI, CCLasso and the implementation of SparCC in SPIEC-EASI. Here PCC and SCC are implemented in R language, and the candidates of associations are selected via p-value. We set p-value at 0.05 for PCC, SCC and CCREPE, and the threshold of correlation for SparCC is 0.1. For each parameter setting, we randomly generate 20 sets of data for evaluation. For all experimental results, it should be noted that we show the mean and variance of evaluation results from the 20 synthetic datasets.

#### Evaluation Metrics

We use three metrics for evaluation:

- **ROC curves:**We plot the ROC curves using two criteria. For PCC, SCC, CCREPE, SparCC and CCLasso, which estimate pairwise correlations, we compare their results with the true correlation matrix*p* with each element being 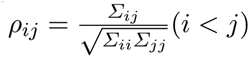. For glasso, SPIEC-EASI and mLDM, which estimate conditional independence, we compare their results with the true precision matrix *Θ*.
- **AUC score:** We compute the area under the ROC curves directly. The AUC scores are calculated by ignoring the sign of edges.
- Δ_1_ **distance:** It is defined as the *L*_1_-distance between the estimated edge weights and the true weights inthe graph. A smaller *Δ*_1_ distance indicates a higher accuracy. Let 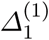 and 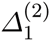 denote the *Δ*_1_ distance for the OTU-OTU and EF-OTU association graphs, respectively. For the pairwise correlation methods, 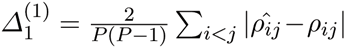, where 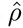 is the estimated value and ρ is the true value. For the conditional independence methods, 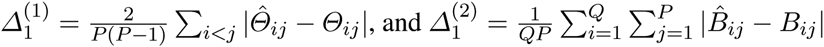.

#### Performance on OTU-OTU Associations

Figure 4 shows the ROC curves of the OTU-OTU association studies for the five different types of graph structures with simulation parameters *P* = 50, *Q* = 5, and *N* = 500. The corresponding AUC scores and *Δ*_1_ distances are summarized in Table 1. From the ROC curves, we can observe that the mLDM model has larger true positive rates than any other methods at small false positive rates. The AUC scores of the mLDM model are generally superior to those of all other state-of-the-art methods on the five different graph structures. In particular, on the Hub structure, the true positive rates of mLDM are significantly higher than those of the other methods. A direct comparison between the mLDM model and the two other methods which estimate conditionally dependent associations without considering the variance of metagenomic data, i.e., glasso and SPIEC-EASI, shows that the mLDM method achieves the highest AUC scores on all five structures. We also observe that the mLDM method has smaller 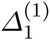 distances than most other methods, suggesting that the mLDM model is able to accurately estimate the weight and sign of conditionally dependent associations. On the cluster graph, the ROC curves of SparCC and CCLasso increase more slowly than those of mLDM at the beginning, but climb higher as the false positive rates become larger. This can be explained by the local density of each standalone cluster in the graph. Under these conditions, mLDM tends to shrink edges with low weights, finally retaining fewer edges than either SparCC or CCLasso. However, we argue that an initial high true positive rate, when the false positive rate is small, is very significant, essentially because a higher ratio of predicted associations will be correct.

**Fig. 4.**
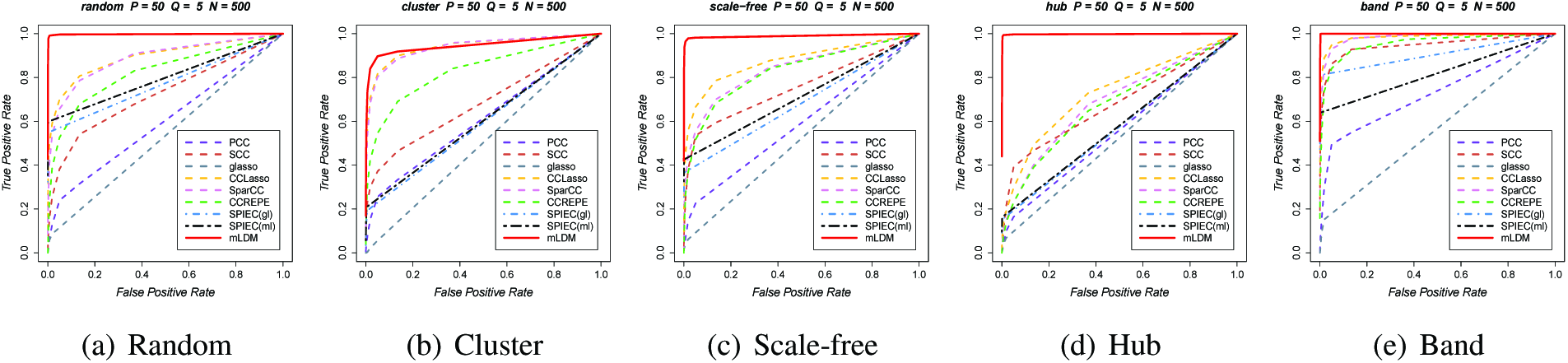
ROC curves of all methods used to discover OTU-OTU associations among microbes when *P* = 50, *Q* = 5, and *N* = 500. These are the average results of 20 simulations with the same parameters.

**Table 1.** *Δ*_1_ distances and AUC scores of OTU-OTU associations with standard deviations *(P* = 50, *Q* = 5,
and *N* = 500).

| Graph<br>Method | Random |  | Cluster |  | Scale-free |  | Hub |  | Band |  |
| --- | --- | --- | --- | --- | --- | --- | --- | --- | --- | --- |
| | $\Delta_1^{(1)}$ | AUC<sup>(1)</sup> | $\Delta_1^{(1)}$ | AUC<sup>(1)</sup> | $\Delta_1^{(1)}$ | AUC<sup>(1)</sup> | $\Delta_1^{(1)}$ | AUC<sup>(1)</sup> | $\Delta_1^{(1)}$ | AUC<sup>(1)</sup> |
| PCC | 0.029 ± 0.0011 | 0.600 ± 0.016 | 0.043 ± 0.010 | 0.610 ± 0.010 | 0.023 ± 0.0010 | 0.590 ± 0.022 | 0.054 ± 0.0006 | 0.559 ± 0.012 | 0.025 ± 0.0022 | 0.732 ± 0.022 |
| SCC | 0.033 ± 0.0032 | 0.728 ± 0.018 | 0.045 ± 0.0035 | 0.684 ± 0.020 | 0.019 ± 0.0012 | 0.758 ± 0.029 | 0.050 ± 0.0017 | 0.684 ± 0.022 | 0.017 ± 0.0022 | 0.950 ± 0.015 |
| CCREPE | 0.028 ± 0.0015 | 0.834 ± 0.019 | 0.039 ± 0.0015 | 0.844 ± 0.017 | 0.022 ± 0.0012 | 0.841 ± 0.025 | 0.050 ± 0.0012 | 0.691 ± 0.025 | 0.025 ± 0.0026 | 0.962 ± 0.008 |
| SparCC | 0.016 ± 0.0005 | 0.899 ± 0.011 | **0.021 ± 0.0010** | 0.945 ± 0.006 | 0.014 ± 0.0005 | 0.854 ± 0.016 | 0.046 ± 0.0012 | 0.709 ± 0.017 | 0.013 ± 0.0005 | 0.985 ± 0.005 |
| CCLasso | 0.020 ± 0.0008 | 0.899 ± 0.016 | 0.026 ± 0.0009 | 0.945 ± 0.007 | 0.017 ± 0.0007 | 0.881 ± 0.023 | 0.046 ± 0.0025 | 0.744 ± 0.061 | 0.016 ± 0.0008 | 0.985 ± 0.007 |
| glasso | 0.021 ± 0.0002 | 0.535 ± 0.005 | 0.038 ± 0.0002 | 0.503 ± 0.005 | 0.014 ± 0.0001 | 0.522 ± 0.008 | 0.017 ± 0.0001 | 0.527 ± 0.012 | 0.023 ± 0.0002 | 0.570 ± 0.024 |
| SPIEC (gl) | 0.018 ± 0.0005 | 0.873 ± 0.050 | 0.037 ± 0.0003 | 0.601 ± 0.034 | 0.012 ± 0.0002 | 0.778 ± 0.050 | 0.017 ± 0.0001 | 0.615 ± 0.051 | 0.018 ± 0.0007 | 0.993 ± 0.008 |
| SPIEC (ml) | - | 0.889 ± 0.041 | - | 0.611 ± 0.024 | - | 0.818 ± 0.068 | - | 0.615 ± 0.051 | - | 0.996 ± 0.005 |
| mLDM | **0.009 ± 0.0009** | **0.998 ± 0.004** | 0.022 ± 0.0014 | **0.949 ± 0.019** | **0.006 ± 0.0006** | **0.990 ± 0.008** | **0.009 ± 0.0001** | **0.998 ± 0.004** | **0.007 ± 0.0009** | **0.999 ± 0.000** |

Figure 6 illustrates the true OTU-OTU association network, and the three networks learned by the three methods with the highest AUC scores (CCLasso, SPIEC, and mLDM), as shown in Table 1. The results visually demonstrate that the association network, as computed by the mLDM model, is closest to the true network and that the mLDM model recovers most of the conditionally dependent associations.

**Fig. 6.**
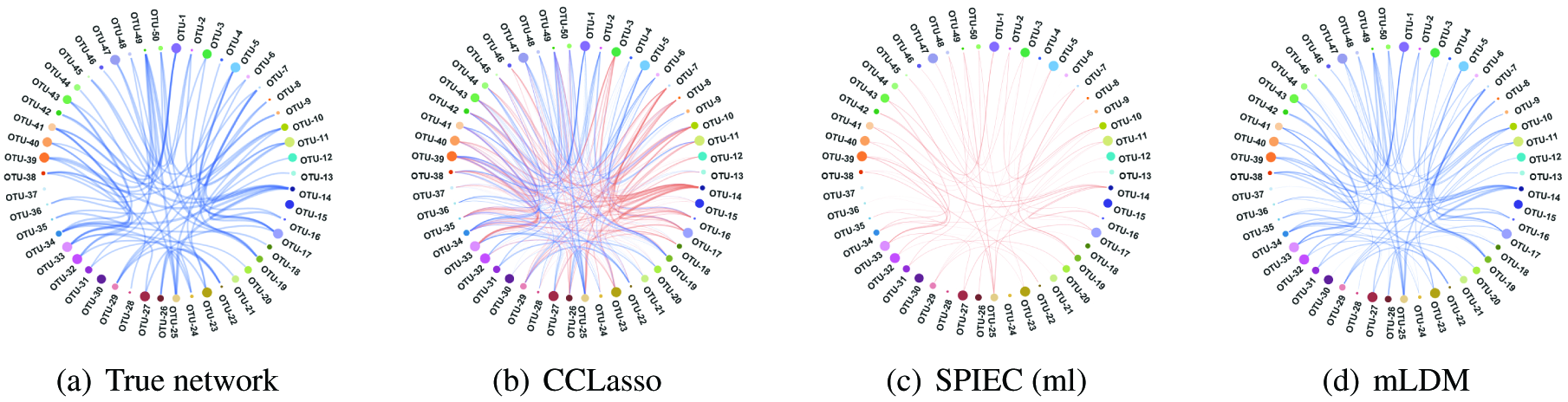
Estimated OTU-OTU associations of three methods (CCLasso, SPIEC and mLDM) and the ground-truth on the ‘Random’ graph (P = 50, Q = 5, and N = 500). The brown and blue curves represent positive and negative and associations, respectively. Thickness of an edge is proportional to the absolute edge weight.

#### Performance on EF-OTU Associations

Figure 5 shows the ROC curves for the estimated associations between environmental factors and OTUs (EF-OTU), where simulation parameters are set as *P* = 50, *Q* = 5, and *N* = 500. The corresponding AUC scores and 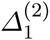 distances are shown in Table 2. Since CCREPE, SparCC, CCLasso and SPIEC do not estimate EF-OTU associations, we compared the mLDM model with PCC, SCC and glasso only. From the ROC curves, we observe that the mLDM model has higher true positive rates and smaller false positive rates than the other four methods. From the AUC scores, we observe that the mLDM model has better performance than the other methods. For 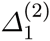 distances, the mLDM model also performs better than the other methods, with the exception of SCC which does slightly better in the Band graph.

**Fig. 5.**
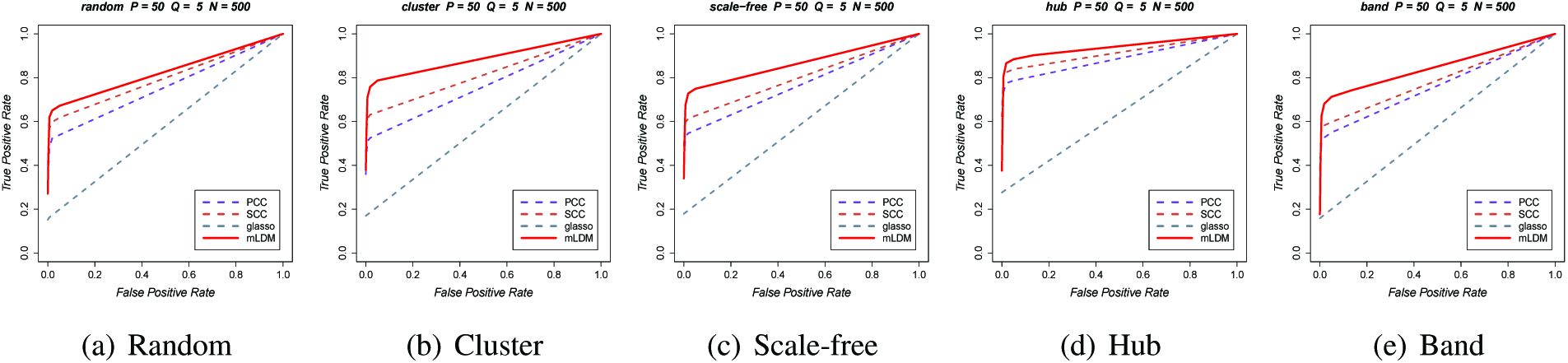
ROC curves of all methods used to discover EF-OTU associations when *P* = 50, *Q* = 5, and *N* = 500. These are the average results of 20 simulations with the same parameters.

**Table 2.** *Δ*_1_ distances and AUC scores of EF-OTU associations with standard deviations *(P* = 50, *Q* = 5, and *N* = 500). Results from other softwares (e.g., CCREPE, SparCC, CCLasso and SPIEC) are omitted here as estimation of EF-OTU is not available.

| Graph<br>Method | Random |  | Cluster |  | Scale-free |  | Hub |  | Band |  |
| --- | --- | --- | --- | --- | --- | --- | --- | --- | --- | --- |
| | $\Delta_1^{(2)}$ | AUC<sup>(2)</sup><sub>P</sub> | $\Delta_1^{(2)}$ | AUC<sup>(2)</sup> | $\Delta_1^{(2)}$ | AUC<sup>(2)</sup> | $\Delta_1^{(2)}$ | AUC<sup>(2)</sup> | $\Delta_1^{(2)}$ | AUC<sup>(2)</sup> |
| PCC | 0.019 ± 0.0013 | 0.774 ± 0.025 | 0.018 ± 0.0011 | 0.790 ± 0.020 | 0.018 ± 0.0014 | 0.813 ± 0.027 | 0.023 ± 0.0019 | 0.896 ± 0.017 | 0.017 ± 0.0019 | 0.778 ± 0.020 |
| SCC | 0.019 ± 0.0021 | 0.804 ± 0.019 | 0.018 ± 0.0021 | 0.833 ± 0.022 | 0.017 ± 0.0020 | 0.840 ± 0.021 | 0.023 ± 0.0030 | 0.916 ± 0.014 | **0.014 ± 0.0010** | 0.792 ± 0.012 |
| glasso | 0.036 ± 0.0004 | 0.649 ± 0.029 | 0.033 ± 0.0004 | 0.645 ± 0.028 | 0.035 ± 0.0004 | 0.654 ± 0.033 | 0.052 ± 0.0005 | 0.735 ± 0.038 | 0.033 ± 0.0003 | 0.641 ± 0.026 |
| mLDM | **0.019 ± 0.0016** | **0.837 ± 0.026** | **0.015 ± 0.0010** | **0.888 ± 0.027** | **0.017 ± 0.0017** | **0.885 ± 0.024** | **0.019 ± 0.0015** | **0.942 ± 0.013** | 0.015 ± 0.0010 | **0.851 ± 0.021** |

Figure 7 illustrates the true EF-OTU association network and the three networks learned by the three methods with the highest AUC scores (PCC, SCC and mLDM), as shown in Table 2. We observe that the networks produced by the mLDM model and SCC are closer to the true network than that by PCC.

**Fig. 7.**
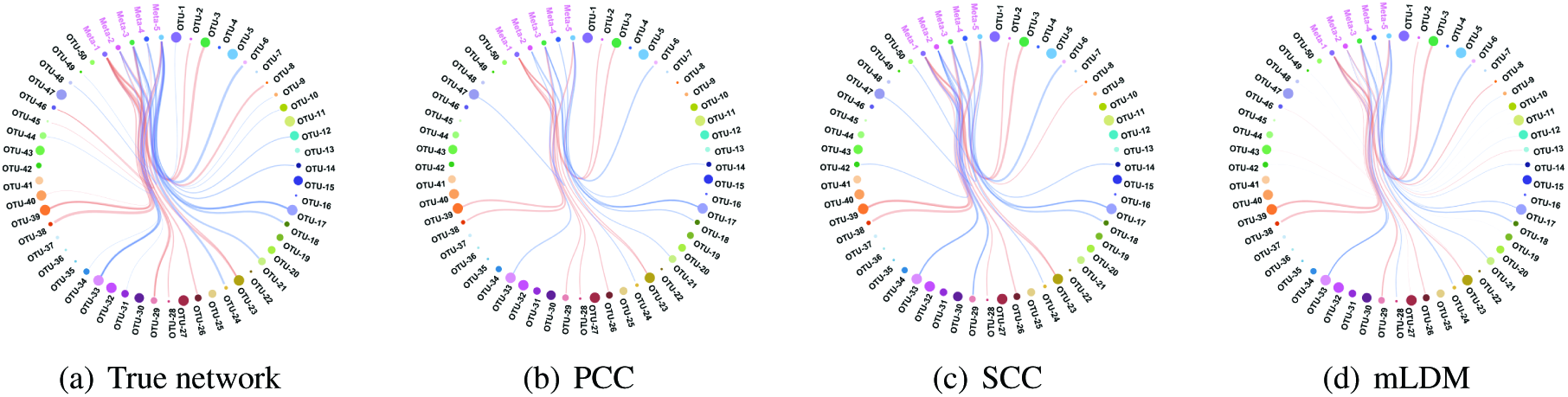
Estimated EF-OTU associations of three methods (PCC, SCC and mLDM) and the ground-truth on the ‘Random’ graph *(P* = 50, *Q* = 5, and *N* = 500). The brown and blue curves represent positive and negative and associations, respectively. Thickness of an edge is proportional to the absolute edge weight.

#### Sensitivity Analysis with Variations of the Sample Numbers

To show the sensitivity of the computational models with respect to different sample sizes, we fixed the number of microbes *P* = 50 and the number of environmental factors *(Q* = 5), and simulated metagenomic sequencing datasets with various sample sizes: *N* = 25, 50,200, and 500. The AUC scores of the estimated OTU-OTU associations by glasso, SPIEC-Easi(gl), SPIEC-Easi(ml) and the mLDM model are plotted in Figure 8. As expected, the AUC scores of all five methods increase when the sample size increases. Among these methods, the mLDM model gives the highest AUC scores on all five graph structures, which again proves that the mLDM model can accurately estimate conditionally dependent associations. The AUC scores of the estimated EF-OTU associations by PCC, SCC, glasso, and the mLDM model are shown in Figure 9 and the AUC scores of the mLDM model are higher than those of PCC, SCC and glasso.

If we combine the performance of OTU-OTU and EF-OTU associations in Figure 8 and 9, we can conclude that the mLDM model outperforms other methods on our synthetic data. The patterns of the ROC curves where N = 25, 50, and 200 are similar to those in Figure 4 and 5 where the number of samples is N = 500, and the initial true positive rate of the mLDM model is better than the others, even though in some case its AUC score is not the highest.

**Fig. 8.**
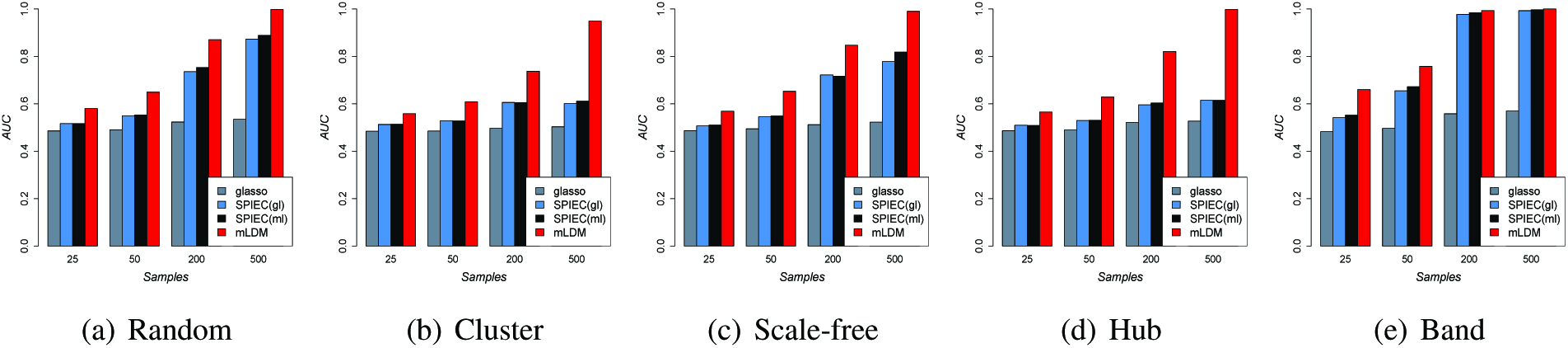
AUC scores of methods which estimate OTU-OTU associations by setting P = 50, Q = 5, and N = 25, 50, 200, and 500.

**Fig. 9.**
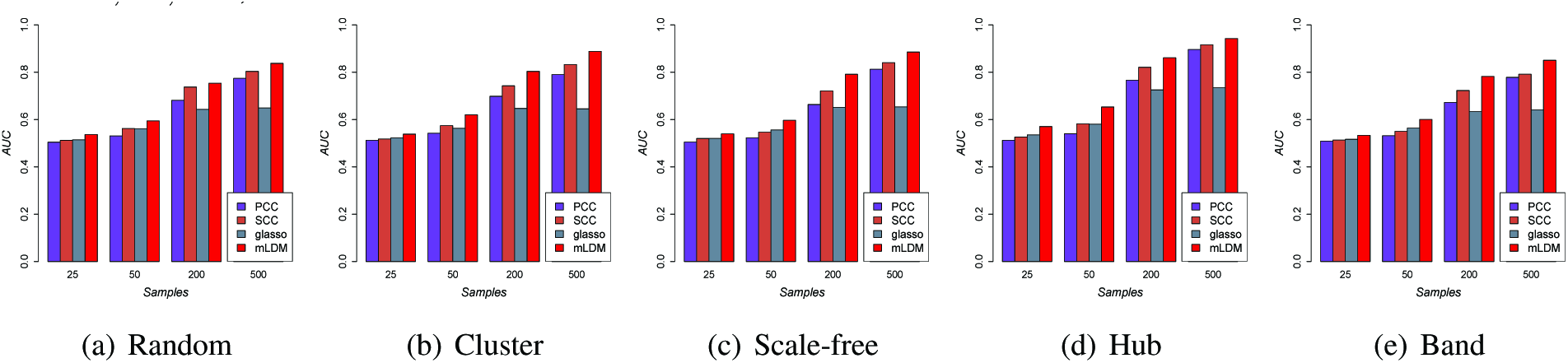
AUC scores of methods used to discover EF-OTU associations (P = 50, Q = 5, and N = 25, 50, 200 and 500.

### 3.2 Tara Oceans Eukaryotic Data

To validate the performance of mLDM on discovering associations from real metagenomic sequencing data, we show the results of mLDM, as well as other eight methods, on Tara Oceans eukaryotic data which were sampled from many stations at different depths over eight oceanic provinces around the world. The eukaryotic abundance profiles were estimated by sequencing and clustering the V9 region of eukaryotic 18s rRNA genes. The OTU table and environmental data, including the known genus-level eukaryotic symbiotic interactions were downloaded from the PANGAEA website^1^ and TARA OCEANS project website ^2^. A total of 91 genus-level mapped eukaryotic symbiotic interactions that consist of both parasitism and mutualism were collected based on the literature [23] and were used to evaluate the effectiveness of all methods.

Samples with missing environmental factor values or with too large or small read counts were removed. OTUs that appear in less than 40% of the samples were omitted. For comparison, we chose OTUs that were involved in known genus-level symbiotic interactions. Eventually we constructed a dataset consisting of 67 OTUs with 28 known genus-level interactions and 17 environmental factors from 221 samples for evaluation.

Given that the known interactions are at the genus-level, and the exact OTU-OTU associations at the species level are unidentified, we evaluated the results at the genus-level. We further specified that a predicted association between two OTUs matches a known genus-level interaction if the two OTUs belong to the two corresponding genera. Considering that the list of known interactions is incomplete, we reported the numbers of matched genus-level associations among the top-N predicted associations (with the highest weights) of all methods, as listed in Table 3.

**Table 3.**
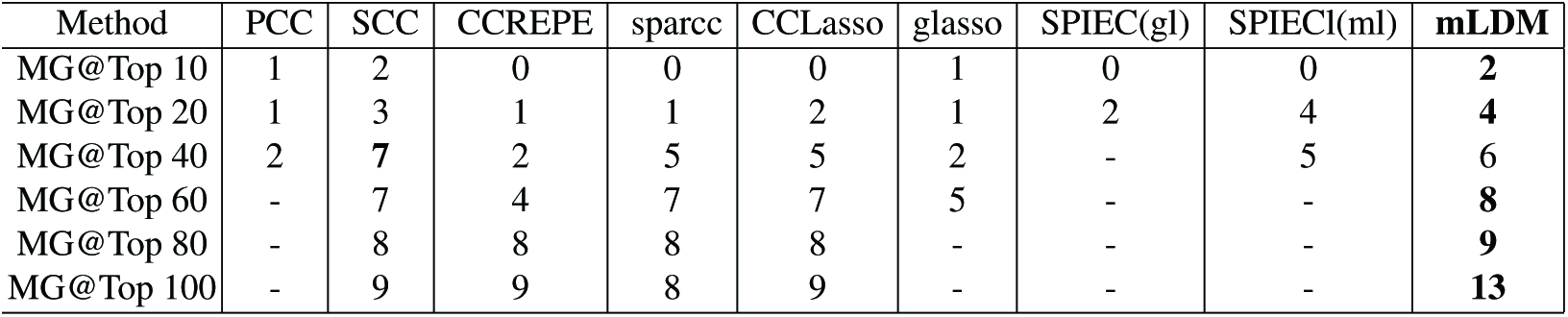
Genus-level associations on TARA Oceans eukaryotic Data. ‘MG@Top N’ is the number of matched known genus-level interactions among top N predicted associations. ‘-’ is shown in the entries where the number of predictions is < N.

| Method | PCC | SCC | CCREPE | sparcc | CCLasso | glasso | SPIEC(gl) | SPIEC(ml) | mLDM |
| --- | --- | --- | --- | --- | --- | --- | --- | --- | --- |
| MG@Top 10 | 1 | 2 | 0 | 0 | 0 | 1 | 0 | 0 | **2** |
| MG@Top 20 | 1 | 3 | 1 | 1 | 2 | 1 | 2 | 4 | **4** |
| MG@Top 40 | 2 | **7** | 2 | 5 | 5 | 2 | - | 5 | **6** |
| MG@Top 60 | - | 7 | 4 | 7 | 7 | 5 | - | - | **8** |
| MG@Top 80 | - | 8 | 8 | 8 | 8 | - | - | - | **9** |
| MG@Top 100 | - | 9 | 9 | 8 | 9 | - | - | - | **13** |

It can be seen that mLDM is superior to other programs in terms of the number of matched associations for 5 out of 6 cases, demonstrating its power of association inference. SCC is competitive with mLDM when N ≤ 40, but its performance decreases as N increases. Both CCLasso and SparCC tend to report a very dense association network, including both true positive associations and a large number of false positive associations, as shown in Figure 11(b) and Figure 11(c). In contrast, mLDM assumes network sparsity and therefore selects associations with higher weights, as shown in Figure 11(a).

**Fig. 11.**
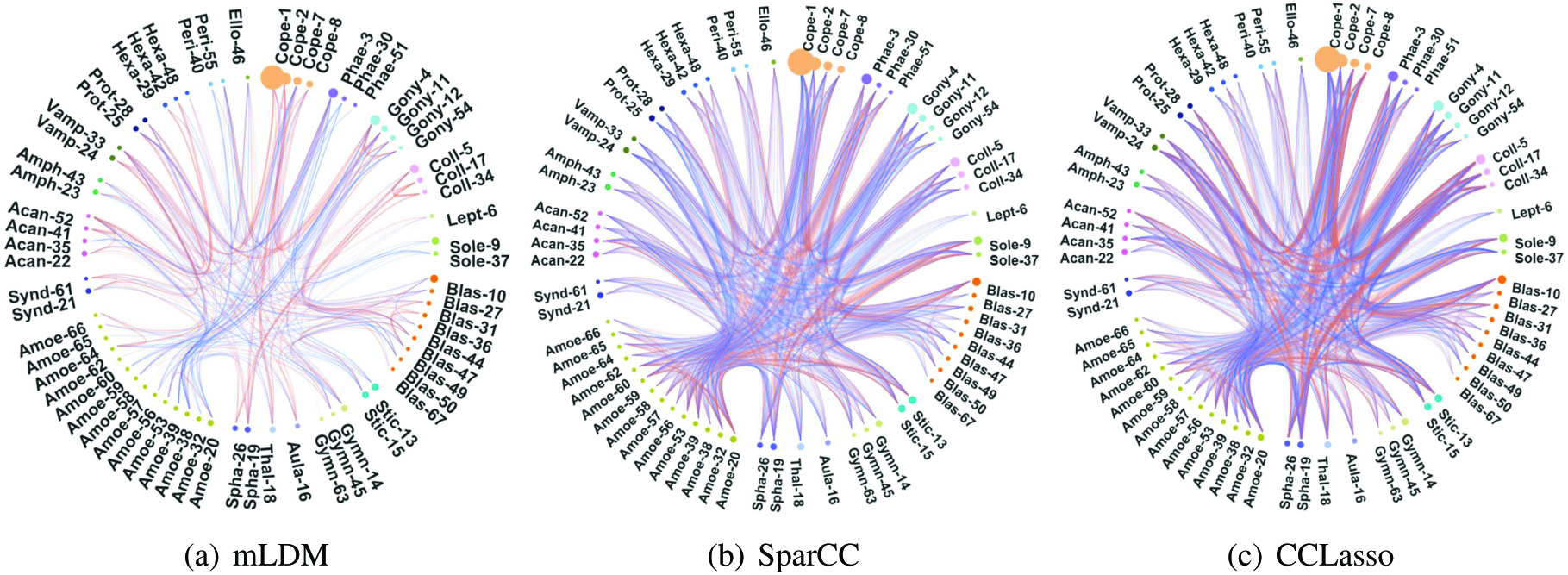
OTU-OTU associations discovered by mLDM, SparCC and CCLasso on TARA Oceans eukaryotic data. The brown and blue curves represent positive and negative associations, respectively. Thickness of edges represents the absolute association weights.

The ground-truth, 28 genus-level symbiotic interactions, as well as the top-40 highest valued genus-level associations discovered by mLDM, are plotted in Figure 10(a) and Figure 10(b), respectively. The strong negative association between the genus *Amoebophrya* and genus *Gonyaulacaceae*, as given by mLDM, implies a parasitism interaction which matches with the known parasitism interactions [10]. Furthermore, the known parasitism interactions between the genera *Amoebophrya* and *Peridiniaceae*, and the genera *Amoebophrya* and *Acanthometra* were also detected by mLDM as having negative associations [24, 8]. However, since the genus V *ampy-rophrya* is positively associated with Amoebophrya, further investigation is required. In addition, mLDM discovered strong positive associations between the genera *Acanthometra* and *Hexaconus*, which are from the same family. We observed similar associations between the genera *Collozoum* and *Sphaerozoum*, which are also from the same family.

**Fig. 10.**
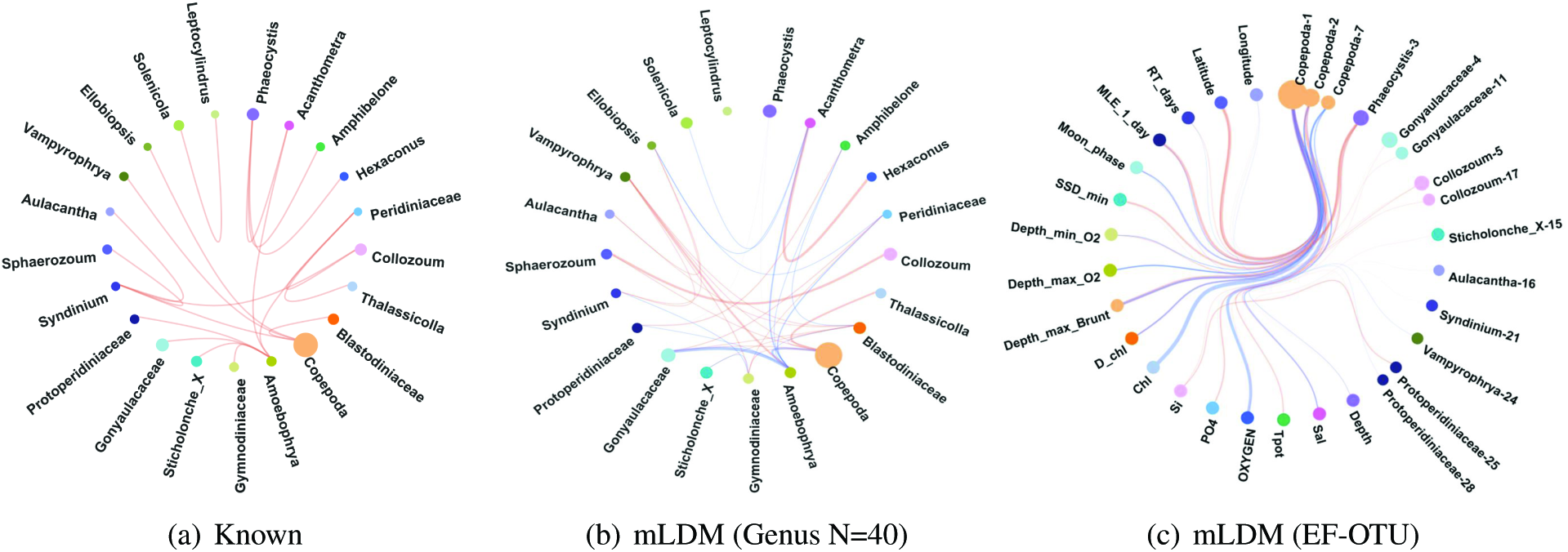
(a) 28 ground-truth genus-level symbiotic interactions where each node represents a genus. Since the signs of the interactions are unknown, we show them in brown for convenience. (b) The genus-level association network discovered by mLDM with its top N = 40 genus-level associations plotted. The brown and blue edges represent positive and negative associations, respectively. Thickness of edges represents the absolute edge weights. (c) Estimated EF-OTU associations discovered by mLDM on 67 OTUs and 17 EFs.

Table 4 lists top 10 predicted OTU-OTU associations (with the largest weights) with relevant literature. For example, two OTUs Hexa-42 and Acan-52 were predicted to be positively associated, which is consistent with the results of a study in the southern California coast by Gilg et al. (2010) [22]. In addition, we found that two associated OTUs Gony-4 and Gony-11 belong to the same genus, and their co-occurrence is consistent with the results of a study of the LSU rDNA sequence data bu Kim et al. (2006) [27].

**Table 4.** Top 10 OTU-OTU and EF-OTU associations on TARA Oceans eukaryotic Data. Associations are sorted in decreasing order according to the weights. The most accurate annotation for every OTU is shown in the bracket. ‘Sign’ records the positive ‘+’ or negative ‘-’ associations, and some relevant studies are listed in ‘Literature’.

| OTU-OTU | Sign | Literature | EF-OTU | Sign | Literature |
| --- | --- | --- | --- | --- | --- |
| Cope-7(Centropages fu.) - Thal-18(Thalassicolla nu.) | + |  | Chl - Cope-1(Corycaeus sp.) | - | [25] |
| Acan-41(Acanthometra sp.) - Acan-52(Acanthometra sp.) | + |  | Latitude - Cope-2(Oithona sp.) | + |  |
| Hexa-42(Hexaconus se.) - Acan-52(Acanthometra sp.) | + | [22] | Depth_max_Brunt - Cope-1(Corycaeus sp.) | + | [26] |
| Coll-17(Collozoum se.) - Coll-34(Collozoum se.) | + |  | Oxygen - Cope-1(Corycaeus sp.) | - |  |
| Gony-4(polygramma) - Gony-11(Alexandrium.01) | + | [27] | MLE_1_day - Phae-3(Phaeocystis) | + |  |
| Vamp-33(Vampyrophrya pe.) - Amoe-62(Amoebophrya sp.) | + |  | Depth_max_Brunt - Cope-7(Centropages fu.) | - |  |
| Amoe-38(Amoebophrya ce.) - Peri-40(foliaceum) | - |  | SSD_min - Phae-3(Phaeocystis) | + |  |
| Coll-5(Collozoum se.) - Coll-34(Collozoum se.) | + |  | D.chl - Cope-2(Oithona sp.) | - | [34] |
| Blas-36(Blastodinium.06) - Blas-49(Blastodinium.05) | + |  | Depth_max_O2 - Cope-1(Corycaeus sp.) | - |  |
| Amoe-20(Amoebophrya) - Amoe-53(Amoebophrya sp.) | + |  | Moon_phase - Cope-7(Centropages fu.) | - | [36] |

In the meantime, we also discovered some interesting EF-OTU associations. Figure 10(c) shows the estimated EF-OTU associations discov-ered by mLDM. The number of the EF-OTU associations is less than that of the OTU-OTU associations, as shown in Figure 11(a), indicating that the environmental factors have direct-affect on only a few OTUs, while the other OTUs are affected via OTU-OTU associations. We further observed that the OTU Cope-1 annotated with the strain *Corycaeus sp.* is negatively associated with oxygen concentration. Almost 95% of all 221 samples of the TARA Oceans dataset are either from the surface waters or from the deep chlorophyll maximum subsurface, whose depths range only from 5.374m to 183.31m. From the samples near the ocean surface, the abundance of *Corycaeus sp.* does not increase linearly with the increase of oxygen, but rather tends to be more abundant when the concentration of oxygen is within a certain range, as plotted in Figure 12.

**Fig. 12.**
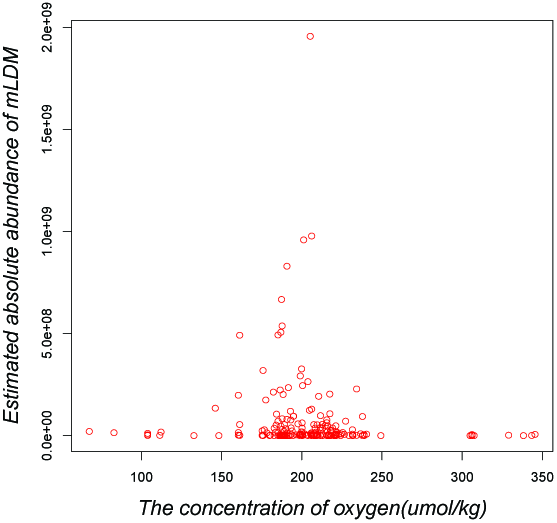
Scatter plot of the concentrations of oxygen and estimated abundances of *Corycaeus sp* by mLDM.

Similarly we show the top 10 predicted EF-OTU associations in Table 4. We found a negatively association between the chlorophyll and Cope-1, which is consistent with the results in a study by Hafferssas and Seridji (2010) [25], in which the association between the chlorophyll and the Copepod structure was found.

Cope-1 is also positively associated with the depth of maximum *Brunt - Väisälä* frequency and the predictability of the depth of maximum *Brunt - Väisälä* frequency to Cope-1 was found by Irigoien et al. (2011) [26]. The relationships between the depth of chlorophyll maximum and Cope-2 and between the moon phase and Cope-7 were also studied in other projects [34, 36].

### 3.3 West English Channel Data

Finally, we applied mLDM to another marine metagenomic sequencing data to infer the underlying OTU-OTU associations and EF-OTU associations. In the marine community, huge amounts of marine microbes exist and play important roles in ocean food chains. However, very little is known about how marine microbes interact with each other or how they are affected by environmental factors. Gilbert et al. [21] studied the dynamic of the marine microbial community in the West English Channel by analyzing high-throughput 16S rRNA data from 2003 to 2008. We downloaded the OTU table from the VAMPS website (https://vamps.mbl.edu/) and employed the mLDM model to analyze the data. Forty-seven samples from position *L4* (50°25.18′N, 4°21.89′W) were selected for association estimation. We extracted 48 OTUs that appear in at least 46 samples. The total abundance of these OTUs exceeds 50% of the total read counts. This dataset has 8 environmental factors, including *temperature, daylength*, as well as *concentrations of salinity, ammonia, chlorophyll, nitrate, phosphate* and *silicate*, which were used to infer EF-OTU associations.

The OTU-OTU association network for the 48 OTUs is shown in Figure 13(a). In general, the number of positive associations (orange edges) among OTUs is more than that of the negative associations (grey edges). The network is clearly dominated by OTUs from *proteobacteria*, which are colored green. The OTU *Alphap* 17, which belongs to the family *Rhodospirillaceae*, plays an important role in the network, as it is the most important hub connected to almost all other OTUs. *Rhodospirillaceae* is known to produce energy through photosynthesis, which is critical to the marine microbial community on the surface of the ocean. Although the OTU *Alphap5* from the genus *Thalassobacter, the* OTU *Alphap2* from the family *SAR11* and the OTU *Alphap* 17 are from the same class, *Alphaproteobacteria*, their associations are different. *Alphap5* and *Alphap* 17 have a strong negative association while *Alphap2* and *Alphap* 17 have a positive association. The OTUs *Gammap* 47 and *Gammap76* are both from the same family, *SAR86*, and have a positive association with OTU *Alphap* 17. It is remarkable that the relative abundance of *Alphap* 17 is low, but still connects with many big OTUs with high relative abundance levels, such as *Alphap1, Alphap2, Gammap76* and *Gammap7*, implying that we should pay more attention to rare OTUs with low abundance in future research.

**Fig. 13.**
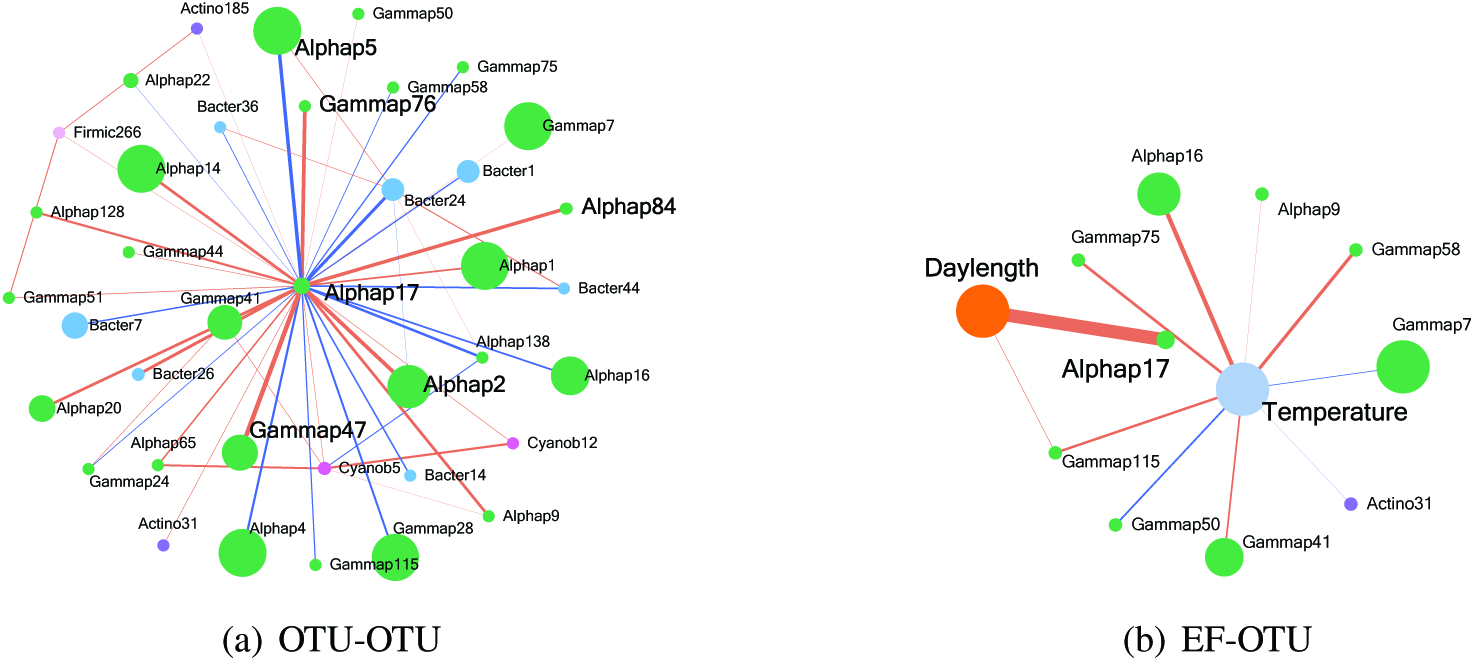
The estimated metagenomic association network of 48 OTUs and 8 EFs. Nodes in the same color belong to the same phylum, and the diameter of each node is proportional to the relative abundance of the OTU. Edges in brown and blue colors denote positive and negative associations, respectively.

Figure 13(b) shows the EF-OTU association network between 8 EFs and 48 OTUs. It can be seen that temperature has the most significant impact on OTUs, especially on the phylum *Proteobacteria.* This is consistent with previous observations. Furthermore, the OTU *Alphap* 17, which connects with many other OTUs, is very strongly and positively associated with day length. This is consistent with the photosynthesis of OTU *Alphap* 17 and further confirms that the photosynthesis of *Alphap* 17 is critical to the whole marine microbial community. In addition, the OTU *Alphap16* from the family *Rhodobacteraceae* has a positive association with temperature. Top 10 OTU-OTU and EF-OTU associations are shown in Table 5. The positive association between temperature and *Alphap16, Gammap58* was reported by Lefort et al. (2013) [30] and Cho and Giovannoni (2004) [12].

**Table 5.**
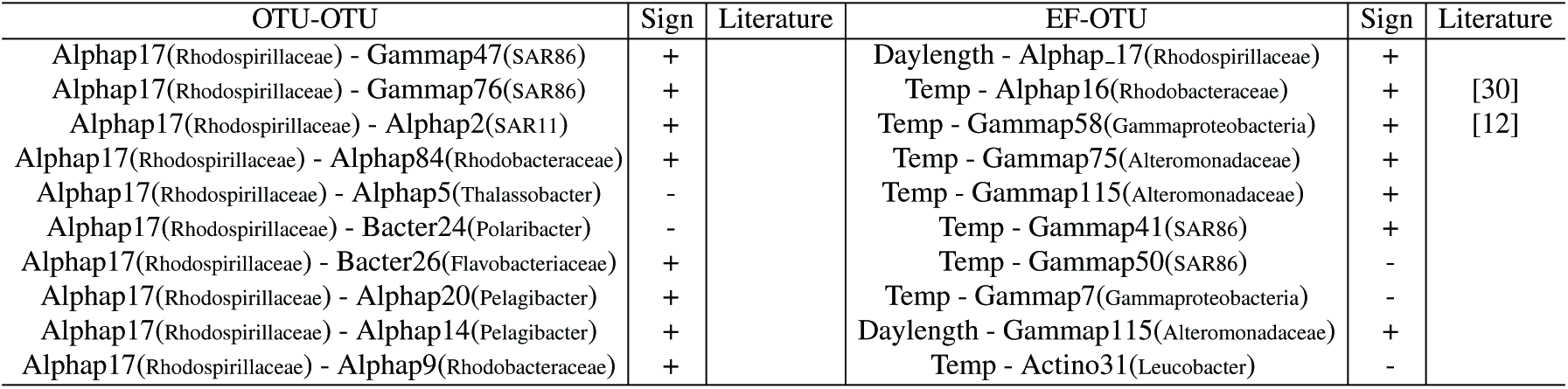
Top 10 OTU-OTU and EF-OTU associations on the West English channel data. Associations are sorted in decreasing order according to the weights. ‘Sign’ records the positive ‘+’ or negative ‘-’ associations. Relevant studies are listed in ‘Literature’.

## 4 Discussions

To discover the underlying associations among microbes from metagenomic samples, we propose a hierarchical Bayesian model, mLDM, with sparsity constraints to discover associations among microbes and between microbes and their environmental factors. The mLDM model can infer both conditionally dependent associations among microbes and direct associations between microbes and environmental factors, by considering both compositional bias and variance of metagenomic data, which have not been studied before. This newly discovered conditionally dependent association provides more insight into the mechanisms underlying a microbial community as it can capture the direct relationship underlying each microbial pair and remove the indirect connection induced from other common factors. The effectiveness of mLDM was verified on the basis of experiments involving both synthetic and real datasets.

It is worth mentioning that LSA was not utilized for performance comparison in the synthetic experiment because it works only for time series data, and our synthetic data are not time-related. In fact, the mechanism of LSA is quite different from other methods mentioned in this paper in that it detects local time-series correlations. Finally, one major limitation of mLDM is its scalability and efficiency, essentially because coordinate descent steps in the hierarchical model consume most of the training time. However, this sacrifice is necessary to gain better performance, which is crucial. For future work, we are interested in developing a scalable mLDM model to analyze extremely large microbial network structures with tens of thousands of microbes by using stochastic gradient descent and parallel computing techniques. For rare OTUs, which only exist in a small fraction of the samples, the lognormal distribution may be not suitable, and other appropriate distributions need to be explored. We are also interested in developing dynamic mLDM models to analyze time series data which is utilized by LSA and learning time-varying network structures.

## Footnotes

1 http://doi.pangaea.de/10.1594/PANGAEA.843018

2 http://www.raeslab.org/companion/ocean-interactome.html

